# Protection of mice against experimental cryptococcosis by synthesized peptides delivered in glucan particles

**DOI:** 10.1101/2021.11.12.468465

**Authors:** Charles A. Specht, E. Jane Homan, Chrono K. Lee, Zhongming Mou, Christina L. Gomez, Maureen M. Hester, Ambily Abraham, Florentina Rus, Gary R. Ostroff, Stuart M. Levitz

## Abstract

The high global burden of cryptococcosis has made development of a protective vaccine a public health priority. We previously demonstrated that a vaccine composed of recombinant *Cryptococcus neoformans* chitin deacetylase 2 (Cda2) delivered in glucan particles (GPs) protects BALB/c and C57BL/6 mice from an otherwise lethal challenge with a highly virulent *C. neoformans* strain. An immunoinformatic analysis of Cda2 revealed a peptide sequence predicted to have strong binding to the MHC Class II (MHC II) H2-IAd allele found in BALB/c mice. BALB/c mice vaccinated with GPs containing a 32 amino acid peptide (Cda2-Pep1) that included this strong binding region were protected from cryptococcosis. Protection was lost with GP-based vaccines containing versions of recombinant Cda2 protein and Cda2-Pep1 with mutations predicted to greatly diminish MHC II binding. Cda2 has homology to the three other *C. neoformans* chitin deacetylases, Cda1, Cda3 and Fpd1, in the high MHC II binding region. GPs loaded with homologous peptides of Cda1, Cda3 and Fpd1 protected BALB/c mice from experimental cryptococcosis, albeit not as robustly as the Cda2-Pep1 vaccine. Finally, seven other peptides were synthesized based on regions in Cda2 predicted to contain promising CD4^+^ T cell epitopes in BALB/c or C57BL/6 mice. While five peptide vaccines significantly protected BALB/c mice, only one protected C57BL/6 mice. Thus, GP-based vaccines containing a single peptide can protect mice against cryptococcosis. However, given the diversity of human MHC II alleles, a peptide-based *Cryptococcus* vaccine for use in humans would be challenging and likely need to contain multiple peptide sequences.

**Importance:** Cryptococcosis, due to infection by fungi of the *Cryptococcus neoformans* species complex, is responsible for substantial morbidity and mortality in immunocompromised persons, particularly those with AIDS. Cryptococcal vaccines are a public health priority yet are not available for human use. We previously demonstrated mice could be protected from experimental cryptococcosis with vaccines composed of recombinant cryptococcal proteins encased in hollow highly purified yeast cell walls (glucan particles). Here, we examined one such protective protein, Cda2, and using bioinformatics, identified a region predicted to stimulate strong T cell responses. A peptide containing this region formulated in glucan particle-based vaccines protected mice as well as the recombinant protein. Other peptide vaccines also protected, including peptides containing sequences from proteins homologous to Cda2. These preclinical mouse studies provide a proof of principle that peptides can be effective as vaccines to protect against cryptococcosis and that bioinformatic approaches can guide peptide selection.

## Introduction

Virtually all cases of cryptococcosis are caused by *C. neoformans* and the closely related species, *C. gattii* (1, 2). The global burden of cryptococcal meningitis has been estimated at 223,100 incident cases per year, with 181,100 deaths (3). The vast majority of patients with cryptococcosis have qualitative or quantitative defects in CD4^+^ T cell function. Cryptococcal meningitis accounts for ~15% of AIDS-related deaths (3). Other immunosuppressed persons are also at high risk; e.g., solid organ transplant recipients have ~1 - 5% lifetime risk of developing cryptococcosis (4). In mouse models of infection, CD4^+^ T cells are also critical for protection, although other arms of the immune system may contribute (5).

Given the public health significance of cryptococcosis, vaccines to protect high risk individuals are a high priority. While heretofore none has reached human clinical trials, promising results have been obtained in animal models (reviewed in (6, 7)). Protection against experimental cryptococcosis can be obtained by immunization with cryptococcal strains missing virulence factors such as capsule, chitosan, sterylglucosidase, and F-box protein (6, 8–11), or genetically engineered to express interferon-γ (12, 13). Whole organism vaccines are relatively easy to manufacture and contain a broad range of antigens. However, they may have difficulty reaching clinical trials due to concerns regarding reactogenicity, autoimmunity and, if administered live, the possibility of causing infection in immunosuppressed persons (14). To circumvent these potential drawbacks, we have focused on identifying candidate antigens, adjuvants, and delivery systems for use in subunit vaccines. We have manufactured vaccines consisting of antigens that are recombinantly expressed in *E. coli* and then encapsulated in glucan particles (GPs) (15, 16). When administered as a prime followed by two boosts, 11 different individual GP-delivered antigens protected BALB/c and/or C57BL/6 mice from pulmonary challenge with the highly virulent KN99α *C. neoformans* strain (15, 16).

Among the most promising of the protective vaccine antigens is chitin deacetylase 2 (Cda2, originally named MP98) (15–17). The immunoreactivity of Cda2 was first demonstrated when it was shown to stimulate a CD4^+^ T cell hybridoma clone created by fusing T cells from immunized mice with a thymoma cell line (17). Subsequently, Weisner *et al*. synthesized a recombinant peptide-MHC II tetramer containing a 13 amino acid peptide from Cda2 (18). Two weeks following pulmonary infection of C57BL/6 mice with the KN99α strain, up to 6.5% of the lung helper T cell population was recognized by the tetramer, thus establishing Cda2 as a major stimulatory antigen. Cda2 belongs to a homologous family which includes Cda1, Cda3, and Fpd1 (also known as Cda4). Cda1, Cda2 and Cda3 have chitin deacetylase activity as shown by their ability to deacetylate cell wall chitin to chitosan in *C. neoformans* (19). Fpd1, which prefers partially deacetylated chitosan as a substrate, may be more properly referred to as a chitosan deacetylase (20). None of the members of the Cda family have significant homology to human proteins (15, 16).

Use of full length recombinant proteins in T cell vaccine studies has the advantage that all epitopes are included in the antigen. However, there is a strong rationale for defining the protective peptides contained within vaccine antigens. First, identifying immunodominant peptide regions of the protein allows elimination of regions of the protein that could drive non-essential, antagonistic, immune suppressive, or autoimmune responses. Second, using synthesized peptides as vaccine antigens minimizes potentially confounding effects of extraneous vector (e.g., *E. coli*)-derived products, such as lipopolysaccharides, lipoproteins, and purification tags. Third, immunoprotective peptides could be combined into a chimeric recombinant protein which would simplify manufacturing and testing of a vaccine in clinical studies (21).

In the present study, we performed an immunoinformatic analysis (16, 22) of Cda2 with the goal of defining CD4^+^ T cell epitopes for use in a *Cryptococcus* vaccine. Peptides within Cda2 were selected based on their predicted binding to the MHC Class II alleles of BALB/c and C57BL/6 mice. Mutated peptides were then created to test the impact of MHC Class II binding on vaccine efficacy

## Results

### Protection with the GP-Cda2 protein vaccine varies as a function of mouse strain

Our published studies (15, 16) and new confirmatory data demonstrate that a GP-based vaccine containing recombinant *E. coli*-derived Cda2 protect BALB/c mice more robustly than C57BL/6 mice (Figure 1). Moreover, DR4 mice, which contain a humanized MHC II allele (DRB1*04:01) on a C57BL/6 genetic background are not significantly protected by the GP-Cda2 vaccine. This led us to hypothesize that the disparities in how well the GP-Cda2 vaccine protected the different mouse strains could be at least partially explained by differences in the MHC II molecules expressed. Our initial focus was on BALB/c mice, given the potent protection mediated by the GP-Cda2 vaccine in that mouse strain.

**Figure 1.**
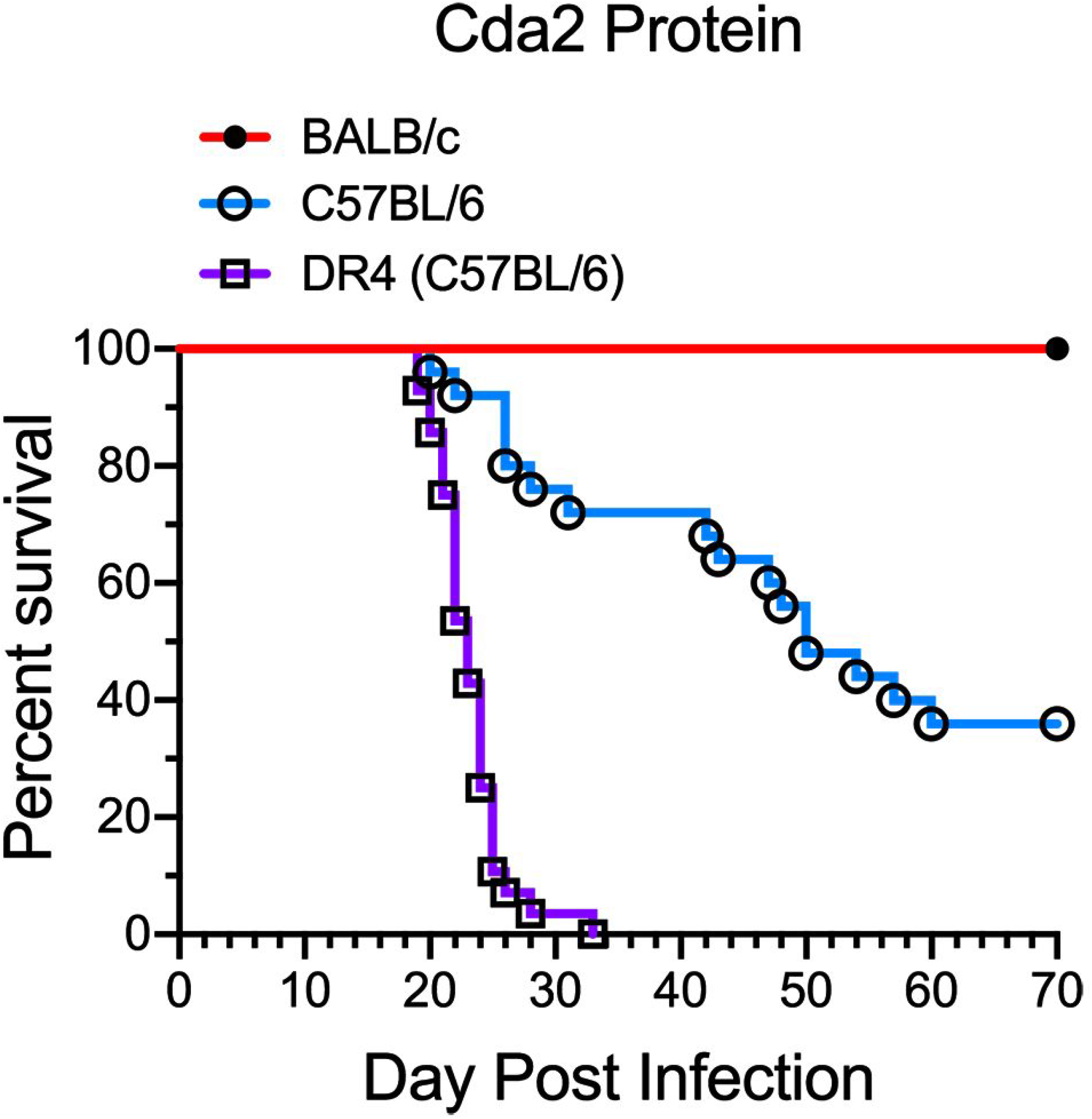
Protection with the GP-Cda2 protein vaccine as a function of inbred mouse strain. BALB/c (n=15), C57BL/6 (n=25), and DR4 (n=28) mice were vaccinated thrice with GP-Cda2 protein and then challenged with *C. neoformans*, as described in *Methods*. Mice were followed daily for survival until day 70 post infection. The figure includes mice previously published (16), as well as confirmatory new experiments. *P≤0.0001* comparing any two groups. Not shown, survival of unvaccinated mice ranged from 20-32 days post infection for each of the mouse strains.

### Mutations in a predicted high binding region of Cda2 result in substantial loss of vaccine-mediated protection in BALB/c mice

We previously identified a region of Cda2 predicted to have 15 amino acid peptides with strong binding to H2-IAd, the MHC II allele expressed by BALB/c [Figure S3 in (16)]. This region also contains the amino acid sequence used to make a tetramer to identify Cda2-specific CD4 T cells following infection of C57BL/6 mice (18). We therefore created mutations in this region of Cda2 spanning amino acids 203-234 of Cda2 (Figure 2A) so that on immunoinformatic analysis, the predicted H2-IAd binding (three positions designated by asterisks in Figure 2B) was greatly diminished or lost entirely. Two such mutated regions, designated M1 and M2 were selected. *E. coli*-derived proteins comprising these mutated sequences were then synthesized, and GP-based vaccines were manufactured. BALB/c mice were vaccinated, challenged via the pulmonary route with the KN99 strain of *Cryptococcus*, and followed for survival over a 70d observation period. Vaccine-mediated protection was robust with recombinant “wild-type” Cda2 protein but was mostly lost when Cda2 proteins (Cda2-M1 and Cda2-M2) containing mutated sequences were used (Figure 2C).

**Figure 2.**
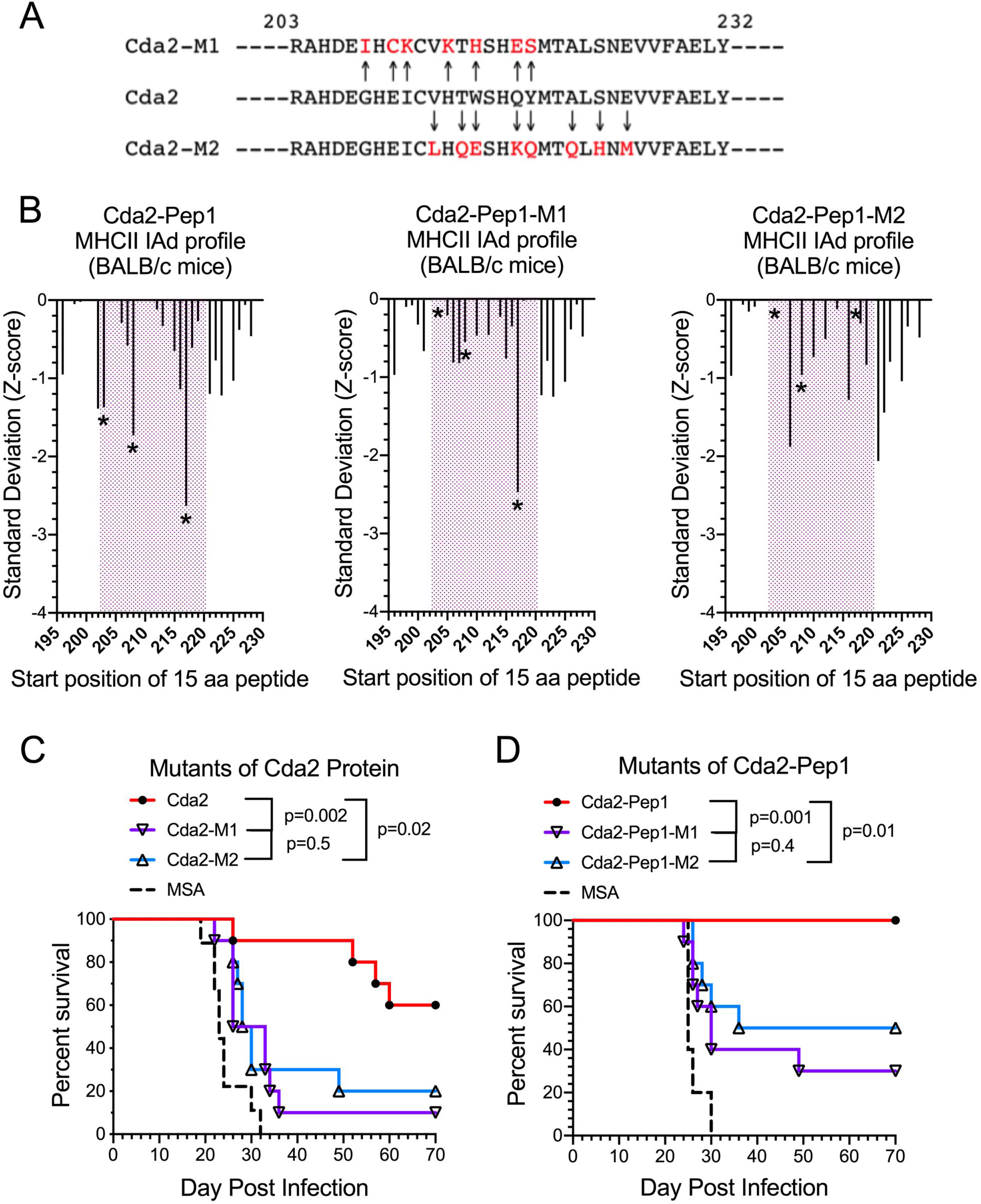
The effect of mutations in a predicted high binding region of Cda2 on GP vaccine-mediated protection in BALB/c mice. A: The sequences of amino acids 203-232 of the wild-type (Cda2-WT) and the Cda2-M1 and Cda2-M2 mutants. Mutated amino acids in Cda2-M1 and Cda2-M2 are indicated by arrows and shown in red. Mutations were designed as in *Methods*. B: Immunoinformatic analysis showing predicted binding of sequential 15 amino acid peptides spanning amino acids 203-232 of WT-Cda2, Cda2-M1 and M2. Predicted binding of each sequential 15mer peptides in CDA2 is shown by index (start) position of the peptide (X axis). The Y axis shows the predicted binding in standard deviation units relative to a mean of zero. A lower Standard Deviation (Z-Score) indicates greater predicted binding, as described in *Methods*. C: Survival studies in mice vaccinated with *E. coli*-expressed Cda2 (wild type), Cda2-M1, and Cda2-M2 protein loaded into GPs. Control mice received GPs containing mouse serum albumin (MSA). The number of mice in each group: Cda2, n=10; Cda2-M1, n=;10 Cda2-M2, n=10; MSA, n=9. D: As in C except synthesized 32 amino acid peptides were loaded into GPs rather than *E. coli*-expressed protein. The number of mice in each group: Cda2-Pep1, n=10; Cda2-M1, n=10; Cda2-M2, n=10; MSA, n=5.

### Protection mediated by peptide vaccines containing the predicted high binding region of Cda2

We next synthesized 32-mer peptides spanning amino acids 203-234 from the predicted high binding region of Cda2, along with the corresponding regions of M1 and M2 mutants. These peptides were named Cda2-Pep1, Cda2-Pep1-M1, and Cda2-Pep1-M2, respectively. Remarkably, mice that received GP-based vaccines containing Cda2-Pep1 were protected from experimental cryptococcosis (Figure 2D). In contrast, protection was diminished, albeit not eliminated, with the vaccines containing Cda2-Pep1-M1 and Cda2-Pep1-M2.

### Protection mediated by vaccines containing peptides homologous to Cda2-Pep1 that are present in other cryptococcal chitin deacetylases

Cda2 has homology to Cda1, Cda3 and Fpd1, including in the predicted MHC II H2-IAd high binding region of Cda2 (Figure 3A). We synthesized 32 amino acid peptides, termed Cda1-Pep1, Cda3-Pep1 and Fdp1-Pep1 based on sequences homologous to Cda2-Pep1. The peptides were loaded into GPs and used to vaccinate mice. Compared with unvaccinated mice, mice vaccinated with any of the GP-Pep1 vaccines were protected against an otherwise lethal pulmonary challenge with *C. neoformans* (Figure 3B). Protection was greatest for vaccines containing Cda2-Pep1, followed by Cda1-Pep1, Cda3-Pep1, and Fpd1-Pep1.

**Figure 3.**
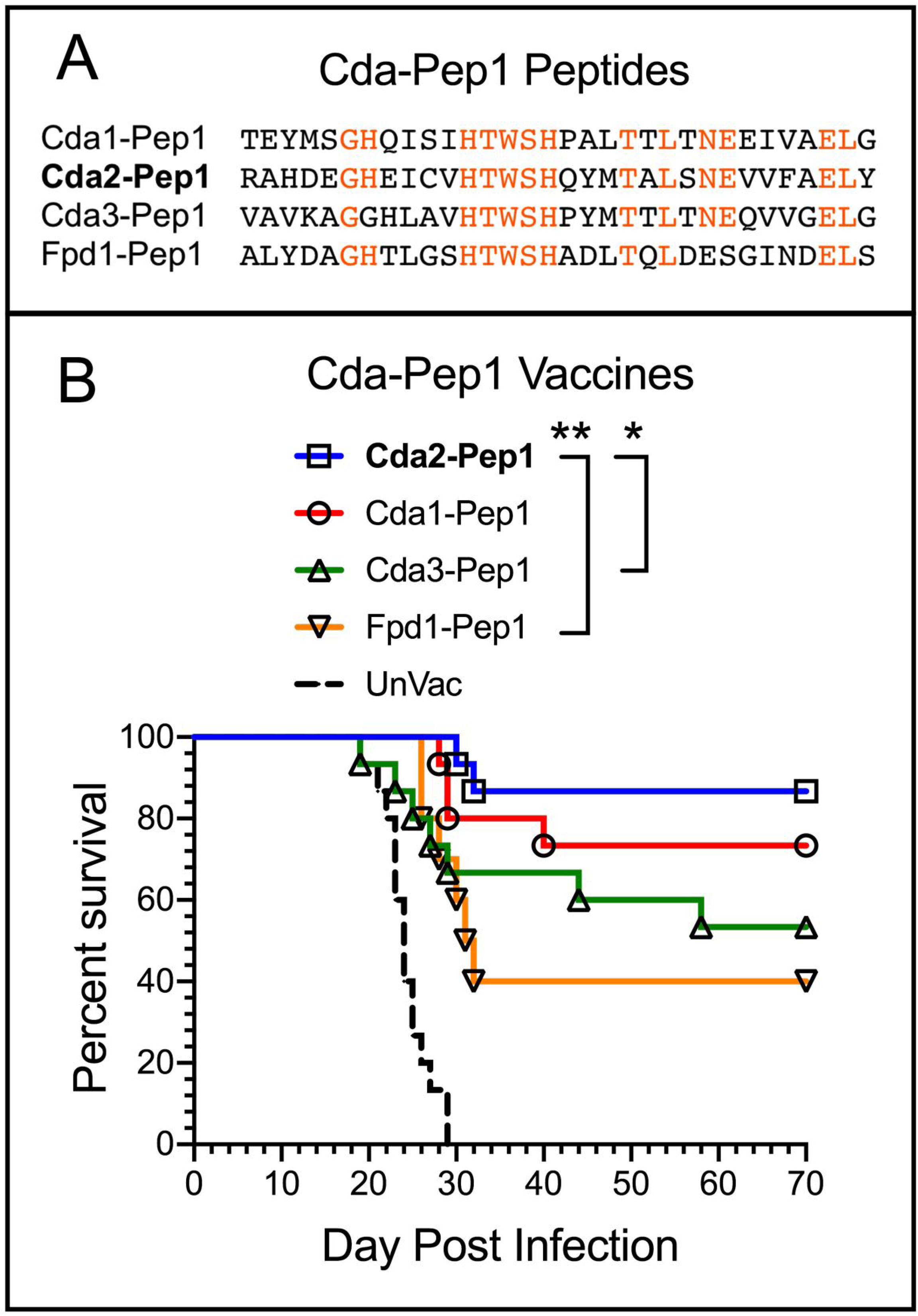
Protection mediated by GP-based vaccines containing peptides synthesized based on cryptococcal chitin deacetylases sequences with homology to the Cda2 predicted high binding region. A. Sequences of the 32 amino acid peptides, Cda1-Pep1, Cda2-Pep1, Cda3-Pep1, and Fpd1-Pep1. Amino acids with identity to the corresponding amino acid in Cda2-Pep1 are shown in red. B. Survival studies in BALB/c mice vaccinated with Cda1-Pep1, Cda2-Pep1, Cda3-Pep1, and Fpd1-Pep1 loaded into GPs and then challenged with *C. neoformans*. UnVac, unvaccinated *C. neoformans*-challenged mice. *, p=0.04 (not significant after Bonferroni correction). **, p=0.009. Not marked on the figure, p<0.0001 comparing UnVac with any of the four vaccinated groups. There were 10 mice in each of the vaccinated groups and 15 mice in the unvaccinated group.

### Protection of BALB/c and C57BL/6 mice mediated by GP-based vaccines containing other peptide sequences of Cda2

In the last set of experiments, we synthesized eight 31-35 amino acid peptides based on sequences in Cda2 (Table 1), loaded them into GPs, and tested the GP-peptide vaccines in BALB/c and C57BL/6 mouse models of cryptococcosis. The eight peptides were chosen to overlap with Cda2-Pep1 or based on regions in Cda2 deemed to contain strong CD4^+^ T cell epitopes based on predicted binding to the H2-IAd allele in BALB/c (Figure 4A) and/or the H2-IAb allele in C57BL/6 mice (Figure 4C). Regarding the GP-peptide vaccines, compared with unvaccinated mice, in BALB/c mice, significant protection was seen in five of the eight vaccines (Figure 4B). In contrast, of the eight peptide-based vaccines, only Cda2-Pep5 significantly protected C57BL/6 mice (Figure 4D). Cda2-Pep5 includes what was predicted to be the strongest H2-IAb in the Cda2 recombinant protein (Figure 4C).

**Table 1.**
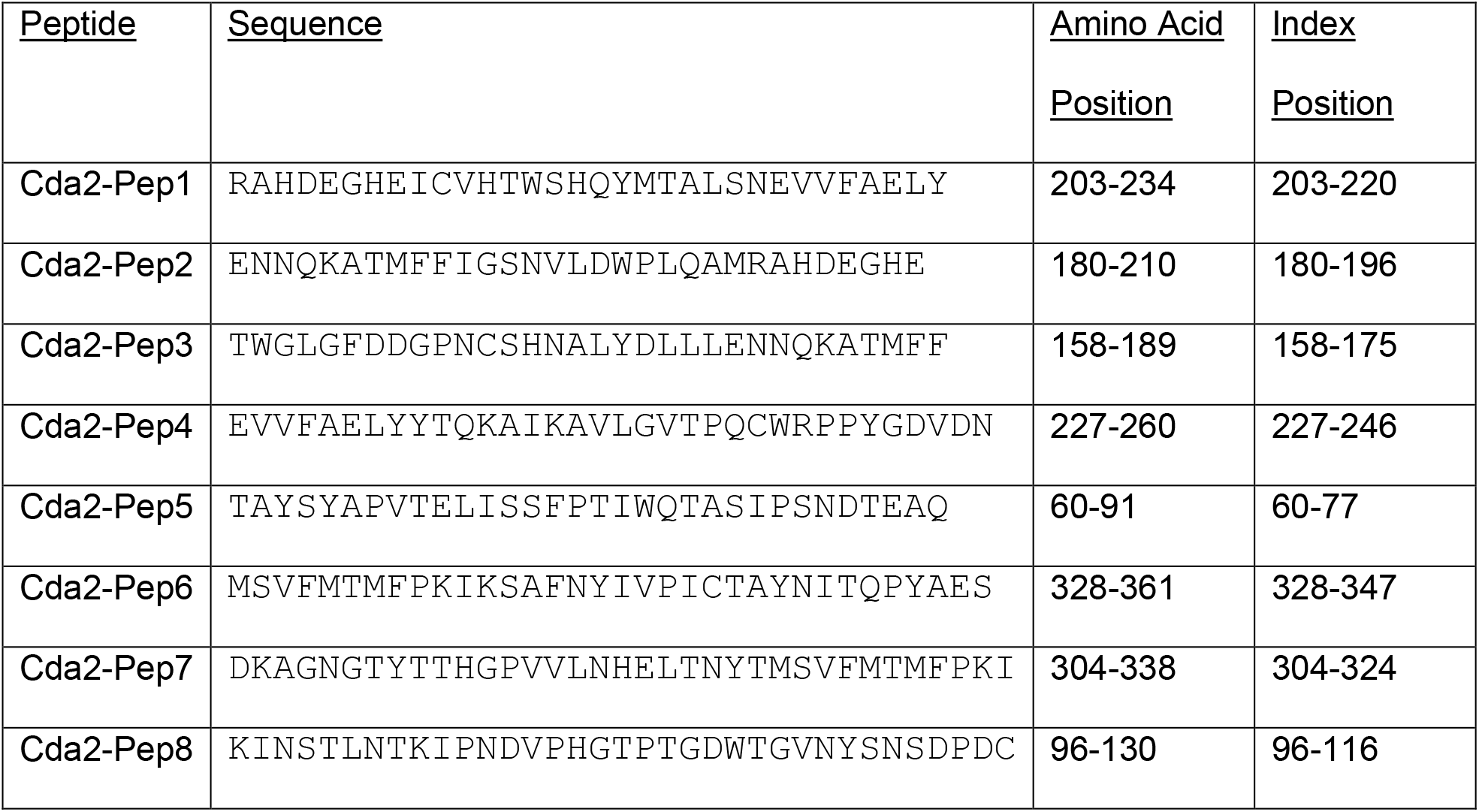
Sequences of Cda2 peptides used in vaccines shown in Figure 4. Protein sequence of Cda2 that was used is from GenBank Accession XP_012049402.1. The index (start) position refers to the first amino acid in a 15 amino acid peptide. The index position is used to identify the vertical bars that depict relative binding of the peptide to a MHCII allele shown in Figures 2B, 4A, and 4C. Thus, in Figure 2B, the vertical bar for Cda2-Pep1 at amino acid 203 refers to the peptide sequence spanning amino acids 203-217.

**Figure 4.**
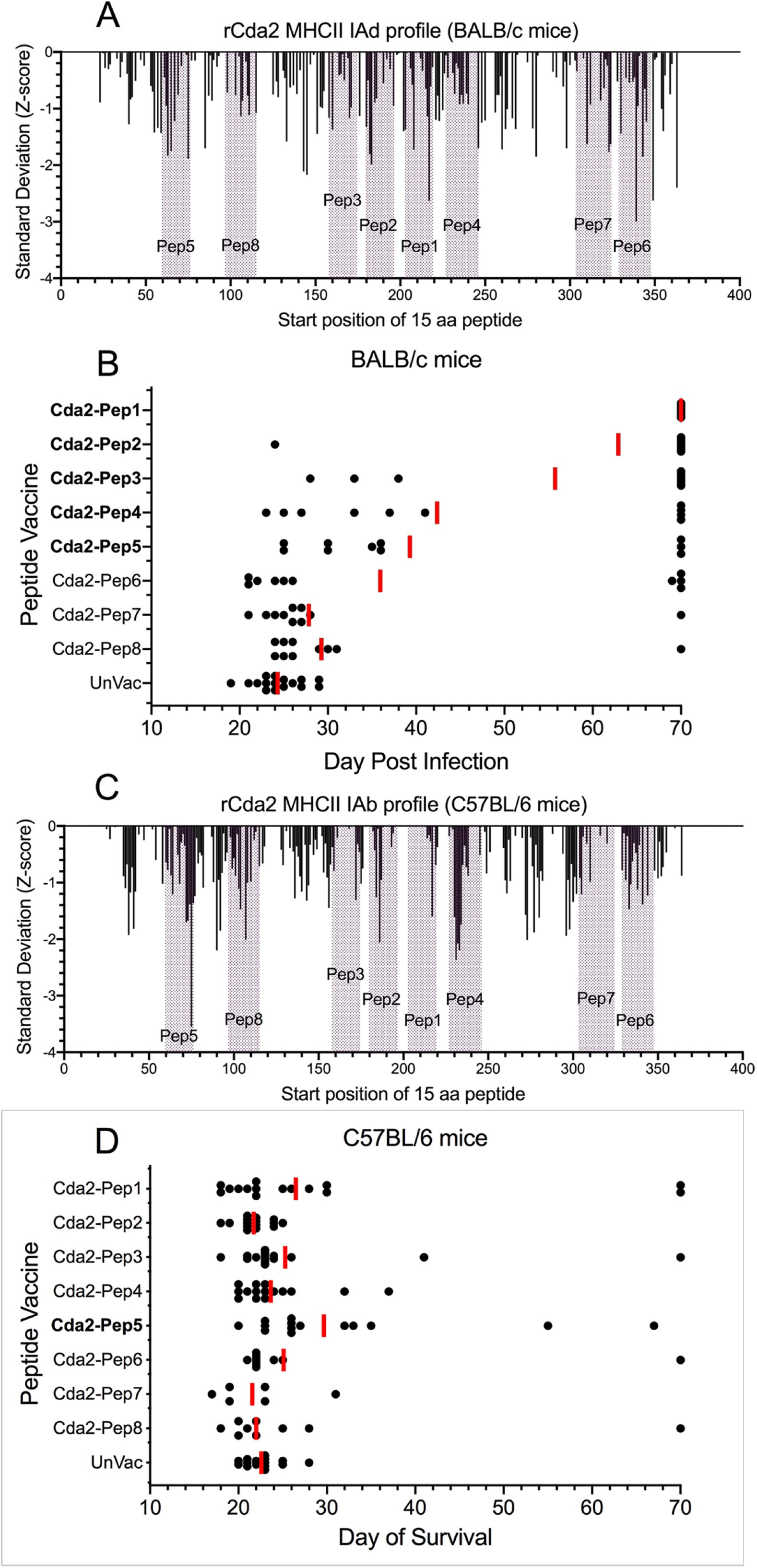
Protection of BALB/c and C57BL/6 mice mediated by GP-based vaccines containing Cda2 peptide sequences. A: Immunoinformatic analysis showing predicted binding to the MHC II IAd allele (present in BALB/c mice) of each sequential 15 amino acid peptide in Cda2 based on the index (start) position of the peptide. The peptides that were tested are colored in purple. The y-axis shows the predicted binding in standard deviation units relative to a mean of zero. A lower Standard Deviation (Z-Score) indicates greater predicted binding, as described in *Methods*. B: BALB/c mice received a prime and two biweekly boosts with the indicated peptide encased in GPs and then challenged with *C. neoformans*, as described in *Methods*. Mice were followed daily for survival until day 70 post infection. Each black dot represents one mouse, shown on the day post infection the mouse succumbed. Each vaccinated group had 10 mice. The experiment was terminated 70 days post infection; survivors were assigned to day 70. The red bars denote the geometric mean survival. UnVac = unvaccinated controls; n=16 mice. Peptide vaccines that afforded significant protection are shown in bold. P<0.0001 for Cda2-Pep1, Cda2-Pep2, Cda2-Pep3, and Cda2-Pep4. P=0.0003 for Cda2-Pep5. C: As in (A), expect the MHC II IAb allele (present in C57BL/6 mice) was interrogated. D: As in (B) except C57BL/6 mice were studied. The number of mice in each group: -Pep1, n=15; -Pep2, n=15; -Pep3, n=15; -Pep4, n=15; -Pep5, n=14; -Pep6, n=10; -Pep7, n=6; -Pep8, n=9; UnVac, n=15. P=0.0005 for Cda2-Pep5.

## Discussion

CD4^+^ T cells are the most critical component of the adaptive protective immune response to naturally acquired cryptococcal infection. A challenge in developing cryptococcal vaccines has been the identification of antigens that induce protective CD4^+^ T cell responses, particularly given the diversity of MHC II in the human population (23). Herein, we performed an in-depth study of an immunodominant protective protein antigen, Cda2, identifying regions of the protein contributing to vaccine-mediated protection in mice.

Our GP-based Cda2-derived peptide vaccines protected BALB/c more robustly than C57BL/6 mice. This is despite the protection afforded both mouse strains by the GP-based vaccine containing the *E. coli*-derived Cda2 protein. Cda2-Pep1, which was only protective in BALB/c mice as part of a vaccine, contains an epitope which is recognized by a sizable fraction of CD4^+^ T cells from infected C57BL/6 mice (18). This emphasizes that immunogenicity does not necessarily result in protection. BALB/c mice are relatively resistant to cryptococcal infection compared with C57BL/6 mice (24, 25). This effect has been attributed in part to a protective Th1 response developing in BALB/c, whereas C57BL/6 mice develop a Th2-biased response (24, 25). While the GP vaccine platform skews towards Th1- and Th17-type responses (26, 27), a response that is broader than just to a single peptide may be required to protect C57BL/6 mice. In addition, the possible contributions of antibody and CD8^+^ T cell immunity must be considered. An alternate but not mutually exclusive possible explanation is the site in Cda2, which comprises the BALB/c MHC II H2-IAd binding site has a functional role that is targeted by the immune response. Of note, Cda2-Pep1 contains two conserved histidine residues required for metal-binding in the catalytic domain of fungal chitin deacetylases and bacterial peptidoglycan deacetylases (28, 29).

DR4 mice are not significantly protected by the GP-Cda2 vaccine. As DR4 mice and C57BL/6 mice each express a different MHC II allele, it is tempting to attribute the disparate protection in the two strains of mice to MHC II binding of processed peptides. However, there is evidence that the CD4-MHC II interaction is impaired in DR4 mice (30, 31). Regardless, the disparate results obtained using different strains of inbred mice emphasize that a successful human T cell vaccine will likely need to contain multiple epitopes given the heterogeneity of the MHC II loci within the human population. Differences in vaccine efficacy comparing mouse strains have also been found using whole organism *Cryptococcus* vaccines (11).

Cda1, Cda2, Cda3, and Fpd1 define a family of homologous chitin deacetylases responsible for deacetylating chitin to chitosan in the cryptococcal cell wall (19). Homology is particularly high in the region contained within Cda2-Pep1, our most protective peptide. GP-based peptide vaccines containing regions in Cda1, Cda2, and Fdp1 homologous to Cda2-Pep1 protected BALB/c mice against cryptococcal challenge. Interestingly, in our previous studies with GP-based vaccines containing recombinant *E. coli*-derived proteins, the same order of protection of BALB/c mice against lethal cryptococcal challenge was observed (i.e., GP-Cda2 was the most protective followed by GP-Cda1, GP-Cda3, and GP-Fpd1) (15, 16). What is unclear though is the extent to which each vaccine elicits cross-protective responses to their homologous family members. For example, is part of the protection mediated by the GP-Cda2-Pep1 vaccine due to recall T cell responses stimulated, for example by Cda1 of *C. neoformans*?

MHC binding predictions focus on the flanking regions of the T cell epitopes, also called the pocket positions. The binding affinity indicates the quantitative relationship of a potential epitope with the cognate T cells, based on the on-off rate of the peptide in the MHC II molecular groove and hence the frequency of interactions between the T cell and epitope. Conversely, the amino acid motifs actually engaging a T cell receptor (the non-pocket residues or T cell-exposed motif) are a qualitative interaction. The T cell-exposed motifs are comprised of the central amino acids of any of the 15 amino acid peptides, typically a discontinuous pentamer comprising positions 2,3,5,7,8 of the central 9 amino acid core (22). While there is considerable homology between the sequences from the four proteins we have examined, there is also sufficient sequence diversity that the T cell-exposed motifs are different between the four proteins. Only Cda2-Pep1 and Cda3-Pep1 share exact identity in just 2 out of the 18 T cell-exposed MHC II motifs present for each of the -Pep1 peptides. Any cross-reactivity among the proteins/peptides would depend mostly on “near neighbor” binding of T cells to similar, but non-identical, motifs (32).

Our previous GP-based subunit vaccine studies used recombinant *E. coli*-expressed proteins as the antigens. Although the proteins were His-tagged and affinity-purified on a nickel column, it is possible that LPS and other bacterial pathogen-associated molecular patterns contaminated the preparations and contributed adjuvanticity to the vaccines. The use of synthesized peptides mitigates this concern. Other advantages of peptide vaccines are reviewed in the Introduction and include elimination of regions of the protein that could drive undesirable responses. However, as seen herein comparing the GP-Cda2-Pep1 vaccine in BALB/c and C57BL/6 mice, a drawback of peptide vaccines is they may be MHC II allele-dependent. A chimeric recombinant protein (21) containing multiple peptide antigens could be designed so that it stimulated Th-dependent responses among a broad range of MHC Class II alleles found in the human population. However, care would need to be taken to avoid creating spurious neoepitopes at peptide junctions and/or dominance hierarchy issues (33, 34).

Our studies add to the growing literature regarding the power of bioinformatics to predict T cell epitopes and inform vaccine development (35, 36). Nevertheless, *in silico* immunoinformatic approaches have limitations because they cannot fully account for posttranslational modifications. Furthermore, modeling has been done only those alleles for which there are adequately large training sets. Relevant to our studies, native *C. neoformans* Cda2 is heavily mannosylated (17, 37, 38), and glycosylation can interfere with antigen processing and presentation (39). While the precise mannosylation sites are not known, each of the peptides we tested contains multiple serines and threonines which are potential sites of O-linked glycosylation. Moreover, peptides Cda2-Pep3, Cda2-Pep5, Cda2-Pep6, Cda2-Pep7, and Cda2-Pep8 contain one or more consensus sequences (Asn-X-Ser or Asn-X-Thr) required for N-linked glycosylation. Ultimately, antigen-specific CD4^+^ T cell-mediated protection is multifactorial; additional determinants may include epitope combinations, binding MHC alleles, T cell repertoires (prior exposure to same or similar epitopes creating a responsive clonal population), and cathepsin and endosomal processing.

Our data serve as a proof of principle that peptide vaccines engineered to stimulate CD4^+^ T cell responses can protect mice against a highly virulent *C. neoformans* strain. The vaccines were adjuvanted and delivered using the GP platform, which biases towards strong Th1 and Th17 responses (26, 40, 41); future studies will be needed to determine whether other adjuvants can be substituted. Given the diversity of MHC II alleles in the human population, a peptide-based vaccine designed for use in humans would likely require multiple peptides. An additional challenge to translate our findings to humans is the impairments in CD4^+^ T cell function present in most individuals at risk for cryptococcosis. Individuals would likely need to be vaccinated when their T cell function was relatively intact, such as early in HIV infection or prior to solid organ transplantation. Moreover, combining a T cell vaccine with one that elicits protective antibodies (42) merits testing.

## Methods

### Reagents, Peptides, and *C. neoformans*

Except where noted, chemical reagents were obtained from Thermo Fisher Scientific. Peptides of >75% purity were synthesized by GenScript and provided as lyophilized material in measured amounts of peptide. Each peptide was analyzed by GenScript for purity using HPLC, Mass spectrometry and nitrogen analysis. Depending on their solubility, peptides were dissolved in water, 50% DMSO, or 100% DMSO. Stock solutions of each peptide were adjusted to 5 mg/ml based on their calculated extinction coefficient (E^0.1%^ at 280 nm; ProtParam tool at Expasy.org) and stored at −80°C. *C. neoformans* serotype A strain KN99α (43) was stored in glycerol stocks at −80°C and grown for in vivo infection studies as described (15, 16). Briefly, following an initial culture on YPD (Difco Yeast Extract, Bacto Peptone, Dextrose) with 2% agar, yeast cells were grown in liquid YPD at 30°C with shaking for 18h. Yeast cells were then harvested by centrifugation, washed with PBS, counted, and suspended in PBS at 2-4×10^5^ cells/ml.

### Recombinant *E. coli*-expressed proteins

National Center for Biotechnology Information file for *C. neoformans* var. grubii H99 strain (taxid:235443) served as the source for cDNA and protein sequences of Cda1 (CNAG_05799), Cda2 (CNAG_01230), Cda3 (CNAG_01239), and Fpd1 (CNAG_06291). cDNAs for these proteins and the mutated versions of Cda2 (Cda2-M1 and Cda2-M2) were synthesized and cloned in pET19b (GenScript) so that the vector-encoded His tag was integrated with the N-terminus of the cDNA. Recombinant protein was made in *E. coli* strain BL21(DE3) (New England BioLabs) using Overnite Express™ TB medium (Novagen) and purified on His·Bind resin (EMD Millipore) in the presence of 6M urea, as described (16). Following elution with imidazole, proteins were dialyzed against 6M urea/20 mM Tris-HCl, pH7.9 and concentrated to 10 mg/ml using Amicon Ultra-15 centrifugal filters (10 kDA cutoff, Merck Millipore). The protein concentration was determined by the bicinchoninic acid (BCA) assay. To assess purity, the recombinant proteins were resolved on SDS-PAGE and stained with Coomassie InstantBlue (Expedeon, Ltd.).

### GP-based vaccines

Recombinant *E. coli*-derived proteins were co-trapped with mouse serum albumin (MSA) complexed with yeast RNA (yRNA) in GPs as described (15, 16). Peptides that were water-soluble were loaded in an identical manner. Peptides in DMSO (5 mg/ml) were loaded by mixing 5 μl peptide per mg hydrated GPs, followed by lyophilization. DMSO (2.5 μl/mg GPs) was added to then “push” the peptides into the core of the GPs, followed by lyophilization. A second “push” with 2.5 μl of water/mg GPs followed by lyophilization completed the loading of peptide. Subsequent steps were the same as was done for protein: MSA was loaded in 0.9% saline and the peptide/MSA inside the GPs were co-trapped with yRNA. Following the yRNA trapping step, peptide vaccines were sonicated to single particles, aliquoted, sonicated again and flash frozen. Protein vaccines were washed three times with saline before sonication. Vaccines were stored at −80°C. A vaccine dose consisted of 100 μl of 200 μg GPs (approximately 10^8^ particles) containing 10 μg of recombinant protein or 5 μg of synthesized peptide and 25 μg of MSA complexed with yRNA in 0.9% sterile saline. A control preparation, designated GP-MSA, contained MSA and yRNA without the antigen.

### Mouse studies

C57BL/6, BALB/c, and Abb Knockout/Transgenic, HLA-DR4 (DR4) mice of both sexes were obtained from Charles River Laboratories, The Jackson Laboratory, and Taconic Biosciences. Mice were bred and housed in a specific pathogen-free environment in the animal facilities at the University of Massachusetts Chan Medical School (UMCMS). All animal procedures were carried out under a protocol approved by the UMCMS Institutional Use and Care of Animals Committee.

The vaccination and infection protocols were as described (15, 16). Briefly, vaccinations were administered subcutaneously three times at biweekly intervals. Mice received their first dose of vaccine when 6-10 weeks old. Two weeks following the last booster, the mice were anesthetized with isoflurane and challenged orotracheally with *C. neoformans* strain KN99α. The inoculum for DR4 and C57BL/6 mice was 1 x 10^4^ CFU while for BALB/c mice it was 2 x 10^4^ CFU. Mice were observed twice daily; humane endpoints prompting euthanasia included ataxia, listlessness, weight loss, and failure to groom. The experiment was terminated on day 70 post-infection, at which time all survivors were euthanized.

### Statistics

Kaplan-Meier survival curves were compared using the Mantel-Cox, log-rank test. The Bonferroni correction was applied in instances where multiple comparisons were made, with a P value of <0.05 considered significant after corrections were made. The software program GraphPad Prism Version 9.2.0 was used for all statistical analyses and to generate graphs.

### Immunoinformatics

The immunoinformatics platform used has been described elsewhere (44, 45). Briefly, the mean and SD of natural log of ic50 MHC II allele binding for each sequential 15 amino acid peptide in the protein is predicted by artificial neural network ensembles using algorithms based on vectors derived from the principal components of the physical and chemical characteristics of each amino acid. Mean predicted binding is then standardized to a zero mean unit variance (normal) distribution within the protein to provide a relative competitive index of predicted binding for each peptide in the protein. This places binding predictions of all MHC alleles on the same scale. This metric is expressed in SD units relative to the mean for that protein. Comparison with other prediction systems indicates a predicted binding affinity of <-1 SD units below the mean is a probable epitope (46). The platform also evaluates cathepsin cleavage probability, and the frequency of any T cell exposed (non-pocket) motif relative to reference databases of the human proteome and bacteria of the gastrointestinal microbiome (47–49).

Alterations in the sequence of Cda2-Pep1 were designed to generate sequences with reduced H2-IAd binding affinity. This was done by generating 50,000 random iterations of Cda2-Pep1, progressively replacing 2-8 designated amino acids, and re-evaluating predicted binding of each constituent 15 amino acid peptide. A subset of peptides was then subjected to closer examination to select M1 and M2, each of which has diminished binding to H2-IAd at positions 203, 208 and/or 217. While the T cell exposed motifs in the region of interest are changed, there were no significant differences in the frequency of the exposed motifs relative to the reference databases, indicating that no obvious changes in T cell precursor frequency for the mutant peptides were created. Figure 2 shows the differences in predicted binding of sequential 15 amino acid peptides within -M1 and -M2 compared to the original Cda2-Pep1 peptide.

## Acknowledgments

We acknowledge grants AI025780 (SML), AI102618 (SML), AI125045 (CAS and SML), and AI072195 (CAS), and Contract Number 75N93019C00064 from the National Institute of Allergy and Infectious Diseases (NIAID), National Institutes of Health, USA.

The funders had no role in study design, data collection and interpretation, or the decision to submit the work for publication.

